# Core Genome Multi Locus Sequence Typing and Single Nucleotide Polymorphism Analysis in the Epidemiology of *Brucella melitensis* Infections

**DOI:** 10.1101/297010

**Authors:** Anna Janowicz, Fabrizio De Massis, Massimo Ancora, Cesare Cammà, Claudio Patavino, Antonio Battisti, Karola Prior, Dag Harmsen, Holger Scholz, Katiuscia Zilli, Lorena Sacchini, Elisabetta Di Giannatale, Giuliano Garofolo

## Abstract

The use of whole genome sequencing (WGS) using next generation sequencing (NGS) technology has become a widely accepted method for microbiology laboratories in the application of molecular typing for outbreak tracing and genomic epidemiology. Several studies demonstrated the usefulness of WGS data analysis through Single Nucleotide Polymorphism (SNP) calling from a reference sequence analysis for *Brucella melitensis*, whereas gene-by-gene comparison through core-genome Multilocus Sequence Typing (cgMLST) has not been explored so far. The current study developed an allele-based method cgMLST and compared its performance to the genome-wide SNP approach and the traditional MLVA on a defined sample collection. The dataset comprised of 37 epidemiologically linked animal cases of brucellosis as well as 71 epidemiologically unrelated human and animal isolates collected in Italy. The cgMLST scheme generated in this study contained 2,687 targets of the *B. melitensis* 16M reference genome (75.4% of the complete genome). We established the potential criteria necessary for inclusion of an isolate into a brucellosis outbreak cluster to be ≤4 loci in the cgMLST and ≤10 in WGS SNP analysis. CgMLST and SNP analysis provided much higher phylogenetic distance resolution than MLVA, particularly for strains belonging to the same lineage thus allowing diverse and unrelated genotypes to be identified with greater confidence. The application of this cgMLST scheme to the characterization of *B. melitensis* strains provided insights into the epidemiology of this pathogen and it is a candidate to be a benchmark tool for outbreak investigations in human and animal brucellosis.

## Introduction

Brucellosis is one of the world’s most widespread zoonoses and it is a leading cause of economic losses in production of domestic ruminants (1, 2). Humans can contract the disease by contact with infected animals or their products, with unpasteurized milk being the most common source of brucellosis in urban populations (3, 4). *Brucella melitensis*, which infects primarily sheep and goats, is the most frequent agent of brucellosis in humans and it leads to the most severe manifestation of the disease (5).

Due to the high public health and economic burden of brucellosis, European countries have applied surveillance, control and eradication programs for many years and most of them have acquired the Officially *Brucella melitensis*-Free (OBF) status. The disease however, still persists in several countries in the Mediterranean Area. In Italy, despite implementation of the brucellosis eradication program for over 50 years, ovine and caprine brucellosis remains endemic in several southern provinces, in Sicily in particular (6). To date, the regions of Italy still not classified as OBF cover approximately 35.5% of the national surface, where 39.9% of all small ruminants are farmed (7, 8).

Efficient and reliable surveillance programs are essential for detection and control of outbreaks and largely depend on collection and access to epidemiological data. Currently, epidemiological investigations rely on the availability of standardized and effective molecular typing methods and analysis tools that allow the public health laboratories to identify and trace the outbreak back to its source.

The identification and the typing of *B. melitensis* is still traditionally performed with the use of biotyping techniques, this methodology however, suffers from inconsistencies and requires handling of the live bacteria. For this reason, PCR-based typing is now commonly used as an alternative to the culture-dependent typing methods (9-12). The results of the classical biotyping schemes categorize *B. melitensis* into three biovars that are of limited epidemiological value as they do not provide sufficient resolution between the isolates.

Moreover, an individual biotype often predominates in particular areas, as seen in Italy where the biovar 3 is the almost exclusively isolated from the local animal populations (13). *B. melitensis* is a highly clonal and monomorphic pathogen which renders its differentiation at strain level very difficult (14). Pattern based techniques such as PFGE and AFLP have been applied in the past but these techniques were not able to differentiate *Brucella* at sub-species level which correlated with low intra and inter laboratory reproducibility (15). In recent years, the typing methods have shifted towards the “genome based” approaches that finally allowed an accurate differentiation between *Brucella* isolates and establishment of a common consensus for the sub-typing schemes of this pathogen (6, 16–18).

To date, the Multi Locus Variable number of tandem repeats Analysis (MLVA) is considered to be the most efficient typing method for *Brucella spp*. Several studies demonstrated MLVA to have a high discriminating resolution in congruence with MLST and sufficient for in depth study of either genome evolution or the outbreak epidemiology (19). According to MLVA scheme, *B. melitensis* population can be divided into West Mediterranean, East Mediterranean and American lineages rather than biovars (20, 21). Moreover, with the development of an international repository, the MVLA data can be stored on web servers and shared between research institutes thus increasing MLVA utility as a tool used for analysis of *Brucella* epidemiology in the world (http://mlva.u-psud.fr/brucella/) (22). However, this typing method has several weaknesses, related both to the nature of variable number tandem repeats (VNTRs) as well as to technical demands of the technique itself (12).

With the advances and the decreased cost of whole genome sequencing (WGS), new methods of pathogen typing, including gene-by-gene comparison using core genome Multilocus Sequence Typing (cgMLST) as well as Single Nucleotide Polymorphism (SNP) calling based on a reference sequence analysis are considered as a suitable and a more informative replacement of the gold standard typing schemes (23–26).

The aims of our study were to develop a cgMLST scheme for *B. melitensis*, to describe WGS analyses using examples of an animal outbreak and sporadic cases of brucellosis in humans and livestock and to evaluate the concordance of the WGS-based results with the data obtained using traditional MLVA methodology.

## MATERIAL AND METHODS

### Study design and *B. melitensis* strains

To evaluate the WGS/NGS approach, we defined two different panels of isolates and we compared the results with MLVA-16. The first panel consisted of 37 epidemiologically linked *B. melitensis* isolated from 21 farms from the provinces of Frosinone, Roma, Isernia and Campobasso in central Italy **Figure 1A**. The second panel comprised of 64 epidemiologically unrelated isolates of *B*. *melitensis* collected in Italy from infected livestock between 2011 and 2017 during National eradication program activities, and seven *B. melitensis* isolated from human cases. The **Figure 1B** shows the geographical origin of the samples.

**Figure 1.**
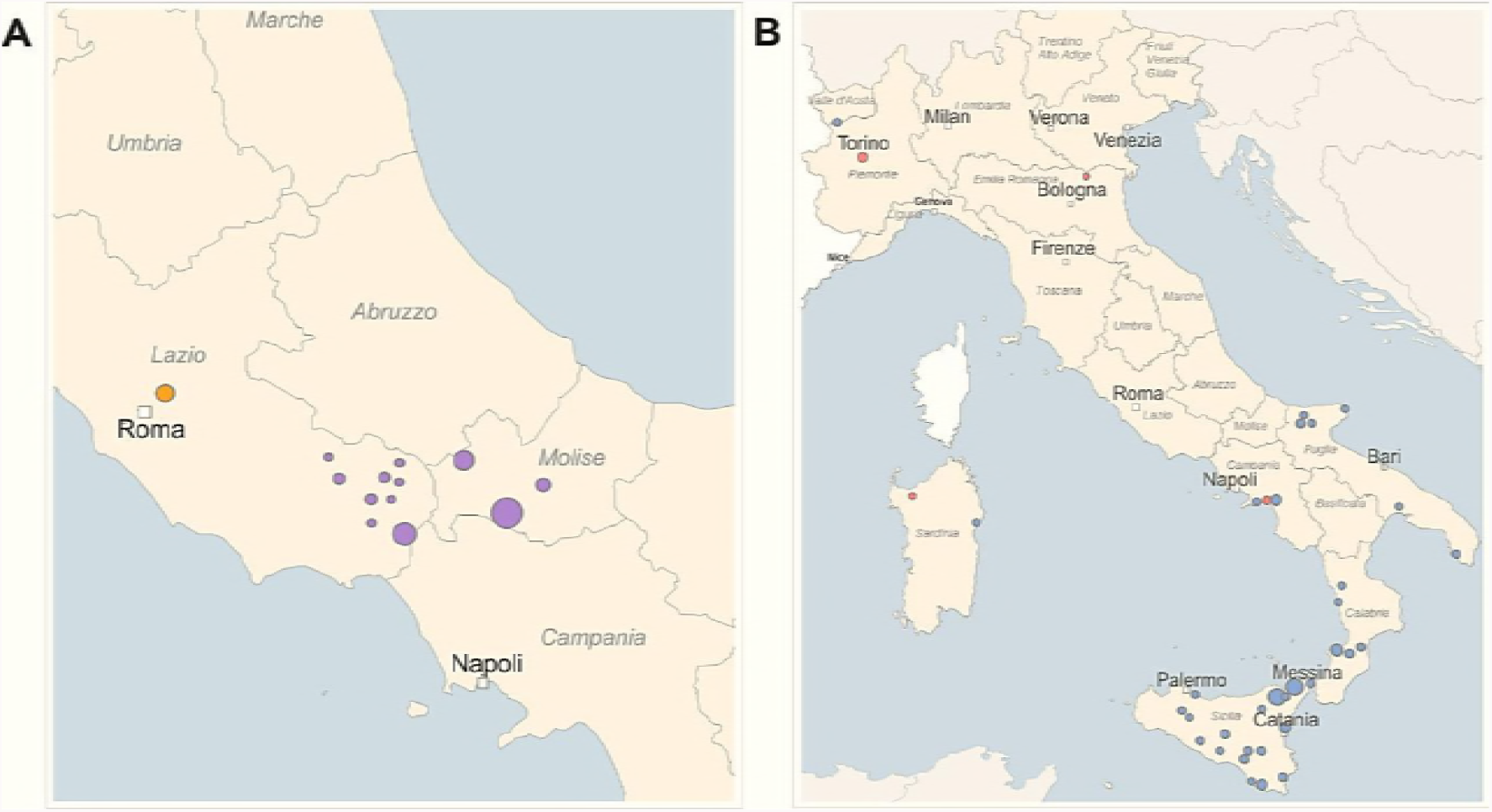
Geographical map for *B. melitensis* cases studied: (A) epidemiologically related isolates and (B) epidemiologically unrelated isolates. Separate epidemiological clusters are marked with different colors respective to the provinces of isolation (purple - Frosinone, Isernia, Campobasso; orange - Roma). (B) The red circles correspond to human isolates and the blue circles to animal isolates.

*B. melitensis* was isolated following the OIE standard bacteriological protocol (27). Briefly, animal samples were collected from lymphatic glands (*i.e*. mandibular, supramammary and genital lymph nodes), spleen, uterus, or udder, whereas human isolates were obtained directly from blood-culture. The isolates were cultured on serum dextrose agar and the phenotype of the colonies was confirmed using a standard Gram stain, catalase, oxidase and urease tests. We assigned the *Brucella* species by PCR, traditional biochemical testing and serotyping. The DNA from the *B. melitensis* strains was extracted using the Maxwell® 16 Tissue DNA Purification Kit, with the Maxwell® 16 Instrument according to the manufacturer’s instructions. All isolates were stored at -80°C, and the associated epidemiological data were recorded as reported in the **Table 1**.

**Table 1.**
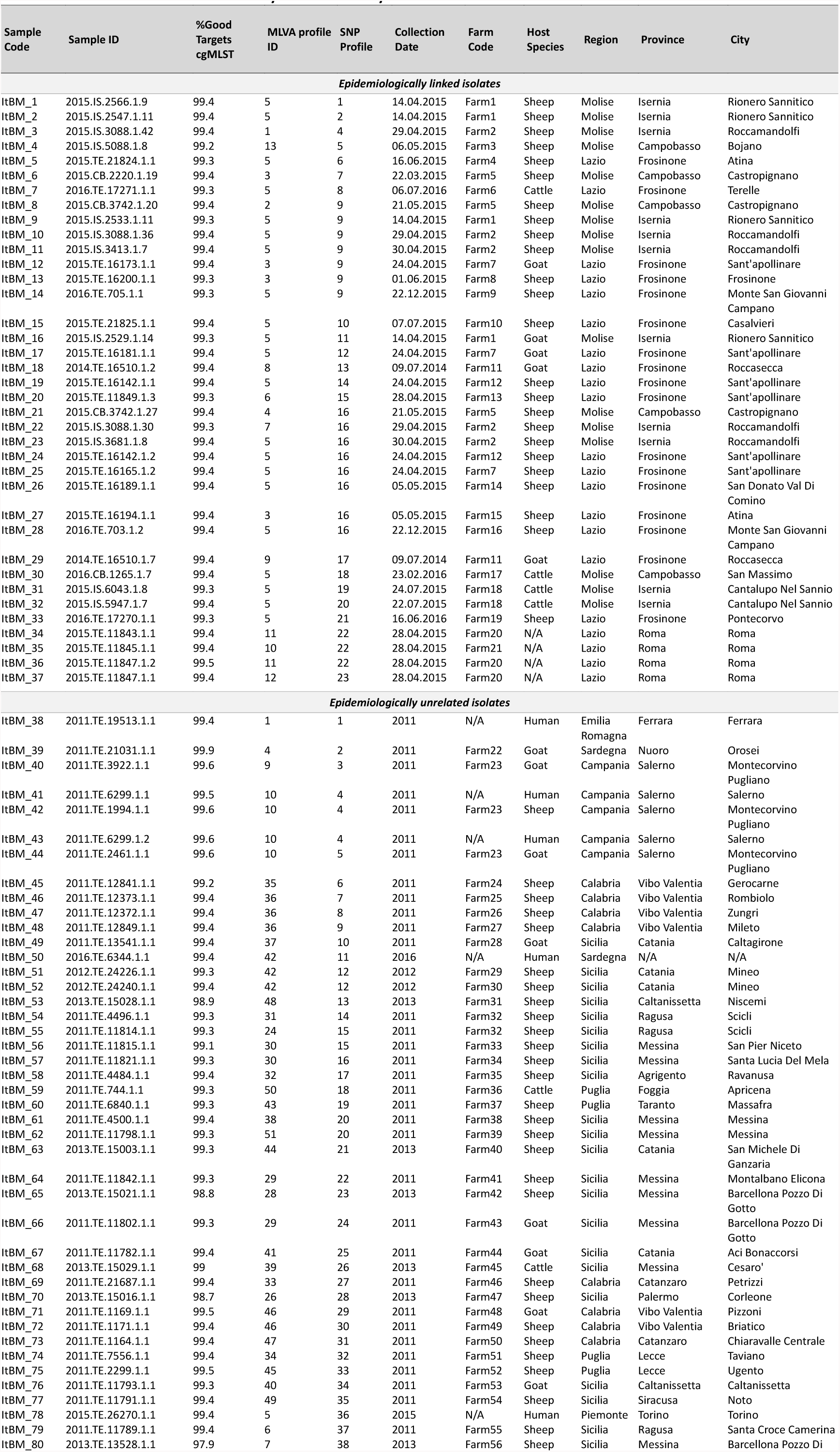
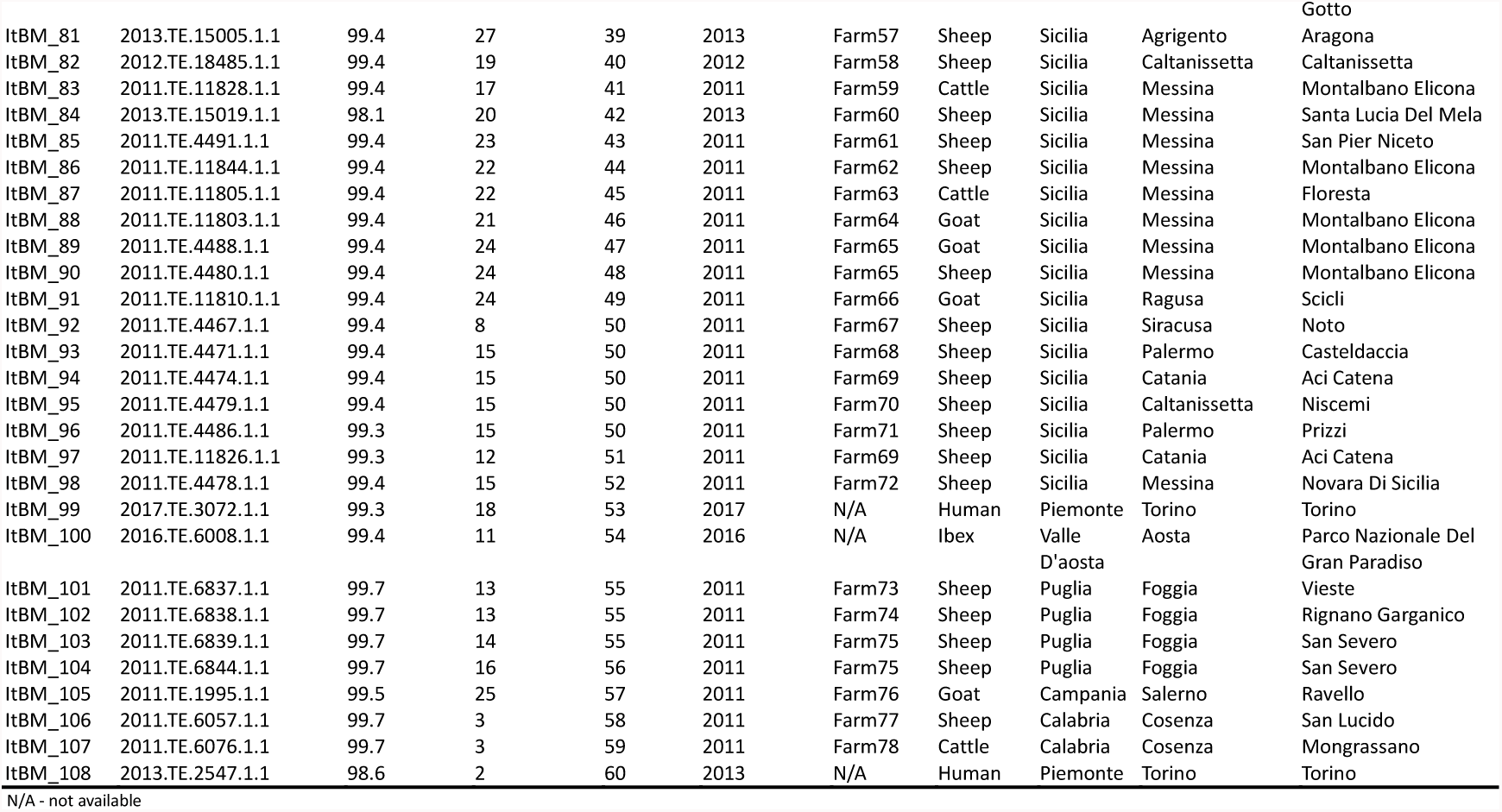
***Brucella melitensis* isolates analyzed in this study**

## MLVA

Samples were genotyped using the MLVA-16 panel described by Le Flèche *et al*. (2006). Briefly, to assign specific allele, DNA extracted from each isolate was amplified by multiplex PCR using primers specific for each of the MLVA 16 loci as described (12). The amplicons were then separated by capillary electrophoresis using ABI 3500 instrument with POP 7 polymer and the allele types were assigned using Genemapper 4.1 (Applied Biosystems, Carlsbad, California, USA).

### Whole Genome Sequencing

Total genomic was DNA quantified with the Qubit fluorometer (QubitTM DNA HS Assay, Life Technologies, Thermo Fisher Scientific Inc.) and library preparation was performed using the Nextera XT Library Prep kit (Illumina Inc., San Diego, CA) or Kapa high/throughput library preparation kit (Kapa Biosystems, Wilmington, MA) according to the manufacturers’ instructions. The libraries were sequenced using Illumina NextSeq 500 platform producing 150bp paired-end reads or Illumina MiSeq producing 250bp paired-end reads. After demultiplexing and removal of the adapters, the reads were trimmed to remove from 5′ end and 3′ end to discard the nucleotides with quality score less than 20. Reads shorted than 70 bp and average Phred mean quality < 24 were automatically discarded. All scaffolds were assembled with SPAdes version 3.11.1 with ‘--careful’ option selected (28, 29).

### cgMLST target definition

To determine the cgMLST gene set, we performed a genome-wide gene-by-gene comparison using the cgMLST target definer (version 1.4) function of the SeqSphere+ software v3.4.1 (Ridom GmbH, Münster, Germany), with default parameters. These parameters comprised the following filters to exclude certain genes of the *B. melitensis* bv.1 str. 16M reference genome (NZ_CP007762.1, NZ_CP007763.1, dated 12 October 2016) from the cgMLST scheme: a “minimum length filter” that discards all genes shorter than 50 bp; a “start codon filter” that discards all genes that contain no start codon at the beginning of the gene; a “stop codon filter” that discards all genes that contain no stop codon, more than one stop codon or if the stop codon is not at the end of the gene; a “homologous gene filter” that discards all genes with fragments that occur in multiple copies within a genome (with identity 90% and more > 100 bp overlap); and a “gene overlap filter” that discards the shorter gene from the cgMLST scheme if the two genes affected overlap > 4 bp. The remaining genes were then used in a pair-wise comparison using BLAST version 2.2.12 (parameters used were: "word size: 11”, “mismatch penalty: 1”, “match reward: 1”, “gap open costs: 5”, and “gap extension costs: 2") with the query chromosomes of representatives for the other two *B. melitensis* biovars (*B. melitensis* bv.2 str. 63/9 NZ_CP007788.1; NZ_CP007789.1 and *B. melitensis* bv.3 str. Ether NZ_CP007761.1; NZ_CP007760.1). All genes of the reference genome that were common in all query genomes with a sequence identity ≥90% and 100% overlap, and with the genome filters “start codon filter”, “stop codon filter” and “stop codon percentage filter” turned on, formed the final cgMLST scheme; this discarded all genes having no start- or stop codon in one of the query genomes as well as genes that had internal stop codons in more than 20% of the query genomes.

### SNP analysis

SNPs were identified using In Silico Genotyper (ISG) version 0.16.10-3 (30). We used default filters to remove SNPs from duplicated regions, minimum quality was set to 30, the minimum allele frequency of 90% and loci not orthologous in all samples. We used ISG pipeline with BWA (31) as the aligner and GATK (32) as the SNP caller. The SNPs were called based on alignment to the reference *Brucella melitensis* bv. 1 str. 16M (GenBank Accession Numbers NC_003317.1; NC_003318.1).

### Clustering analyses

MLVA allelic profiles and SNP matrix data were analyzed using goeBURST algorithm implemented in PHYLOViZ version 2.0 (33). The Minimum Spanning Trees (MST) were created using default software settings. The cgMLST profiles were assigned using *B. melitensis* task template in Ridom SeqSphere+ Software 4.1.1 (Ridom GmbH, Münster Germany) (34). MSTs were created by comparison of cgMLST targets, pairwise ignoring missing values with distance labels representing the number of diverse alleles.

### Accession numbers

All generated reads were submitted to National Center for Biotechnology Information (https://www.ncbi.nlm.nih.gov/) under the bio-project accession number PRJNA448825 (https://www.ncbi.nlm.nih.gov/bioproject/PRJNA448825).

## RESULTS

### Epidemiologically linked *B. melitensis*

The outbreak-related isolates were detected in 21 different farms in three Italian provinces, over a period of one year and a half. The culture-positive samples belonged to 37 animals that were investigated as a part of the within- and among-farm epidemiological investigation (**Figure 1, Table 1)**.

The MLVA revealed the presence of 13 different genotypes divided into 2 groups formed by single locus variants and one double locus variant (**Figure 2A**). The minimum spanning tree (MST) showed that the groups were split by mutations in the three hyper variable loci bruce04, bruce09 and bruce16. One group included three genotypes in four isolates collected from farms located in the province of Rome, whereas in the remaining 34 strains from Isernia, Campobasso and Frosinone provinces we identified ten distinct genotypes. We generated a cgMLST scheme comprising 2,687 targets based on the *B. melitensis* 16M reference genome (75.4% of the complete genome). The cgMLST clustering divided the isolates into two different genetic complexes grouping the two farms from Rome (Complex 2) provinces separately from the remaining nineteen farms (Complex 1). The genetic division measured with the cgMLST panel was by 170 different loci **(Figure 2B)**. The analysis using the “*B. melitensis*” panel found one prevalent genotype that was similar across the provinces of Frosinone, Campobasso and Isernia and was found in ten of the tested farms.

**Figure 2.**
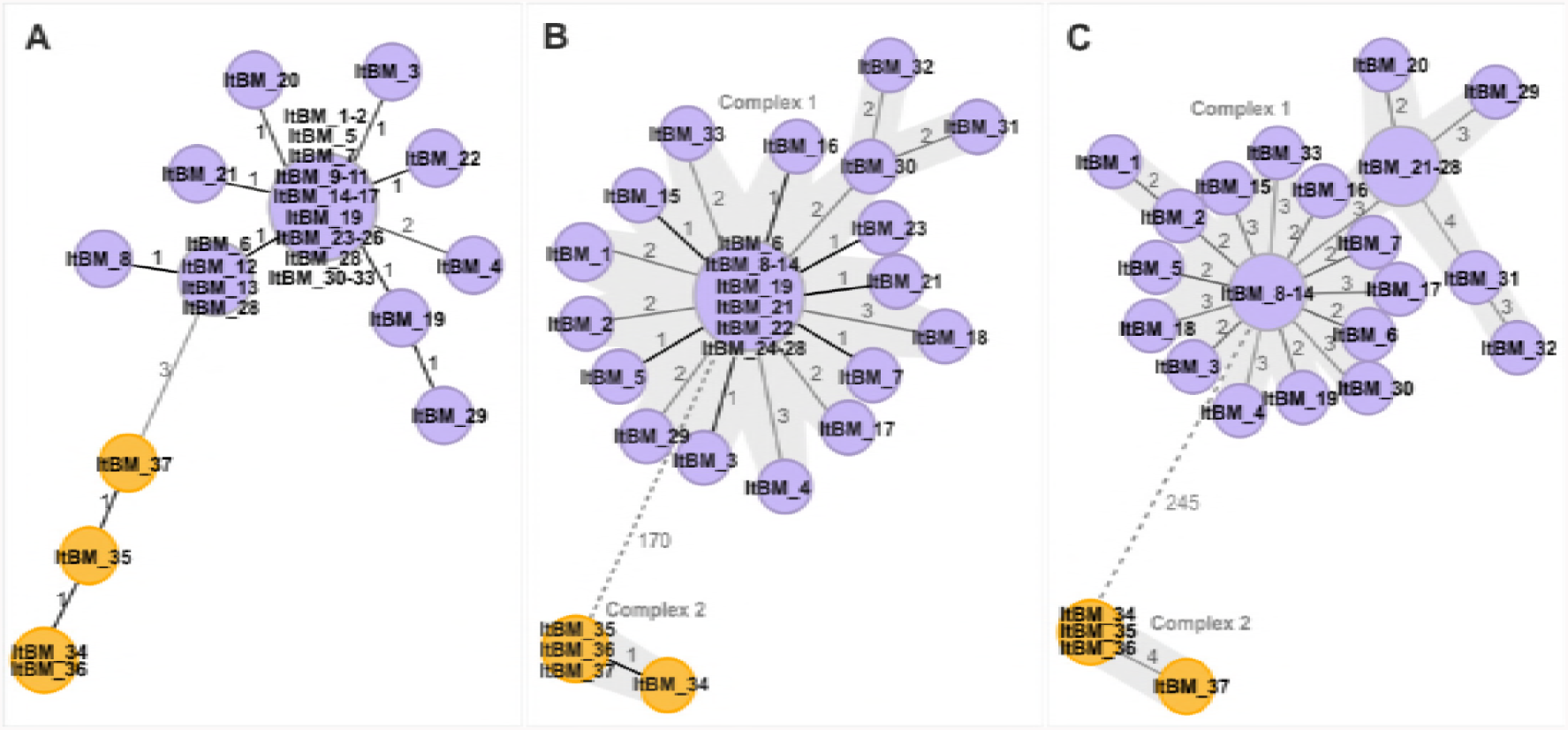
Minimum spanning trees (MST) generated for 37 epidemiologically related isolates. Separate epidemiological clusters are marked with different colors indicating the provinces of isolation (purple - Frosinone, Isernia, Campobasso; orange - Roma). (A) MST based on *B. melitensis* MLVA16 typing. The distance labels correspond to the number of discriminating alleles. (B) MST generated using gene-by-gene approach. CgMLST profiles were assigned using *B. melitensis* task template with 2687 target genes. The MST was created by cgMLST target comparison, pairwise ignoring missing values with distance representing the number of diverse alleles. Separate clusters are highlighted. (C) MST based on single nucleotide polymorphism (SNP) analysis using *Brucella melitensis* strain 16M as a reference. The distance labels correspond to the number of discriminating SNPs between neighboring genotypes.

Seventeen isolates from the Complex 1 shared identical core genome profile and the genetic variation between the isolates in this cluster was not greater than three loci and only one locus in the second cluster. Moreover, removing 52 columns from the analysis where any value was missing, decreased the distances between the nodes even further, and classified all samples from Rome as identical (not shown). A within-farm genetic variation was also observed. In eight farms, samples from more than one animal were analyzed, and several polymorphisms were observed between the isolates originating in the same location, demonstrating that the isolates acquired mutations during the outbreak. (Data not shown).

The SNP analysis from ISG pipeline identified 3,390 SNPs that mapped to 4% of the *B. melitensis* 16M ref strain (NC_003317.1; NC_003318.1), of which 3,146 were classified as clean unique variants and included in the further analysis. The tree split the samples into two genetic clusters with a distance of 245 SNPs between them (**Figure 2C)**.

All the analyses above were in agreement with the epidemiological investigation that established the presence of two distinct and independent introductions of infected livestock.

### Epidemiologically Unrelated *B. melitensis*

The minimum spanning tree (MST) calculated using the MLVA-16 typing results showed a distance between neighboring profiles not exceeding 9 variable loci (**Figure 3**). Fifty-one MVLA profiles were assigned to the 71 strains. Eleven profiles were shared by more than one isolate, which, with exception of one human isolate, corresponded to the samples originating from the same geographical location (**Table 1**).

**Figure 3.**
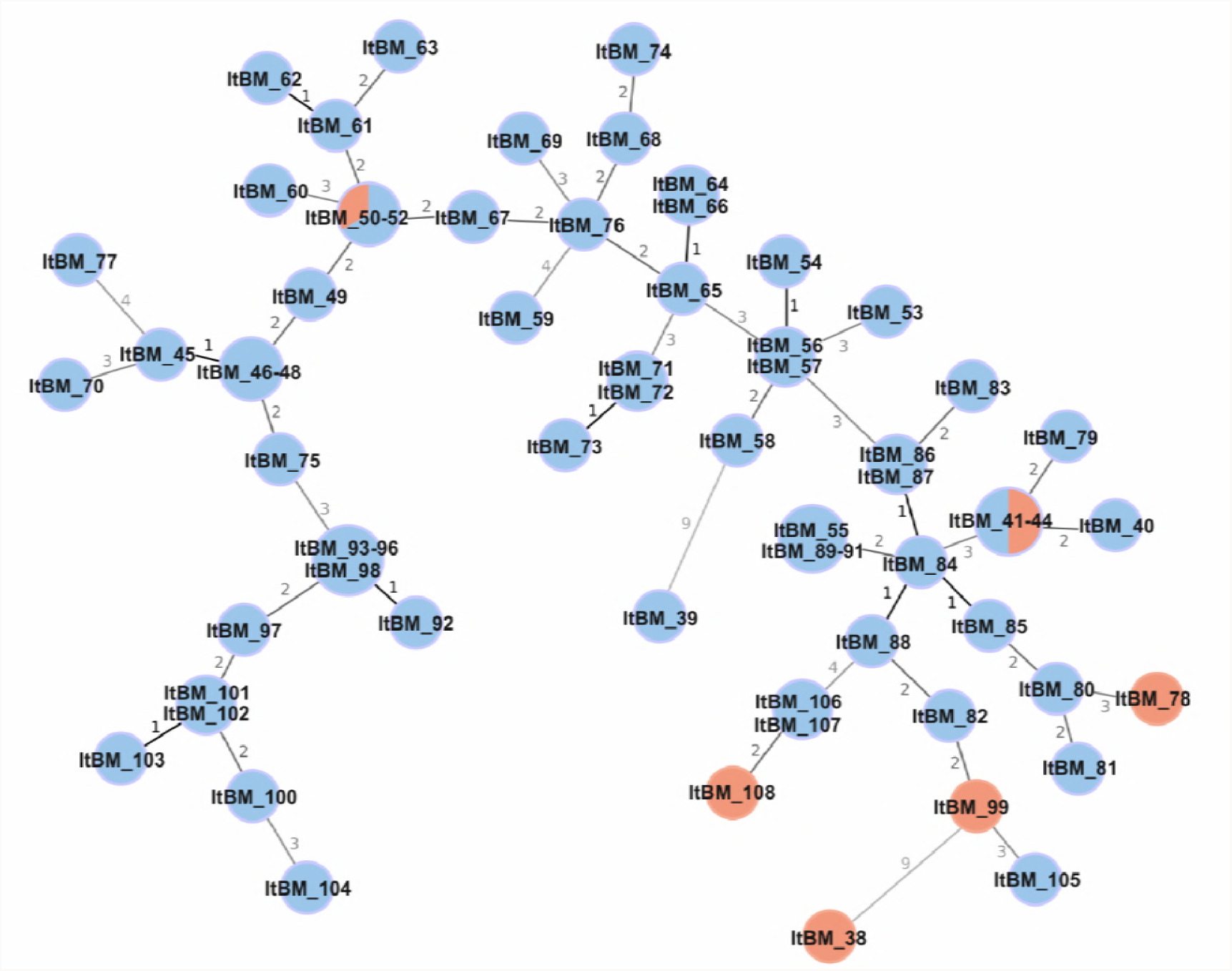
Minimum spanning tree (MST) based on *B. melitensis* MLVA-16 typing results generated for 71 samples from epidemiologically unrelated isolates. The tree was generated using goeBURST algorithm in PHYLOViZ software. The distance labels correspond to the number of discriminating alleles. The red nodes correspond to human isolates and the blue nodes to animal isolates.

MVLA profiles tend to be conserved between epidemiologically linked strains, therefore, the strains from an outbreak are likely to have a similar MVLA profile. Three MLVA profiles, 10 (samples ItBM_41- ItBM_44), 15 (samples ItBM_93 - ItBM_96 and ItBM_98) and 24 (ItBM_55, ItBM_89 - ItBM_91) were identified in more than three strains suggesting close relatedness of samples within these profiles. Samples of profile 10 originated in 2011 in the province of Salerno (~ 4,923 m^2^), which explains their similarity. Isolates from the same year and various provinces of Sicily were found within profile 15, and profile 24 contained four samples from Ragusa and Messina provinces, two of which were collected from the same animal. Apart from MLVA profile 15, which had a neighbor located within one allele of distance (profile 8), the other two profiles were separated from the neighboring ones by at least two alleles.

MLVA typing from the surveillance set allowed identification of two clear outliers. Samples ItBM_38 and ItBM_39 (profiles 1 and 4) showed a distance of 9 alleles from the nearest *B. melitensis* neighboring strain and no relatedness to one another.

According to our MVLA data, only two out of six human cases could be linked to a specific animal source analyzed in our study. Samples ItBM_41 and ItBM_43 isolated from a patient in the city of Salerno (Campania), shared the same MVLA profile as two *B. melitensis* animal isolates from a farm in the municipality of Montecorvino Pugliano from Salerno province (samples ItBM_42 and ItBM_44), all collected in 2011. Human isolate ItBM_50 and two animal isolates (ItBM_51 and ItBM_52) were assigned MVLA profile 42, but interestingly, ItBM_50 was isolated 4 years later than the animal strains. According to this typing method, the same MVLA type had probably circulated in Sicily for at least 4 years prior the human infection occurred. The other three human samples did not show sufficient relatedness to any of the animal isolates to reliably trace the source of infection. The number of variable loci, in these cases, ranged from 2 to 9 in relation to the closest neighboring MVLA profile.

To increase the discriminatory power of the investigation, we analyzed 71 assemblies using cgMLST scheme. All analyzed genome assemblies exceeded 97.9 % of good targets (with a mean of 99.4% good targets). MST tree was built with 55 nodes in total with branch distances between 1 and 1237 genes. Isolates ItBM_38 and ItBM_39 were clear outliers, separated from the closest neighbor by 1,237 alleles, and 1,113 loci from one another **(Figure 4A)**. These isolates were also identified as outliers by the MLVA analysis described above. It is worth to note that these strains were assigned by MLVA to the East Mediterranean and American lineages respectively, whereas all the Italian isolates were a part of the West Mediterranean lineage. Interestingly, cgMLST analysis flagged 3 targets that were uniquely present in these two genotypes but not in any Italian isolates included in this study and therefore they could potentially be used to differentiate West Mediterranean lineage. Upon further inspection, these targets contained 4 extra codons or were missing 4 codons in Italian samples compared with the reference and were therefore classified as failed (threshold was set as default +/- 3 codons).

**Figure 4.**
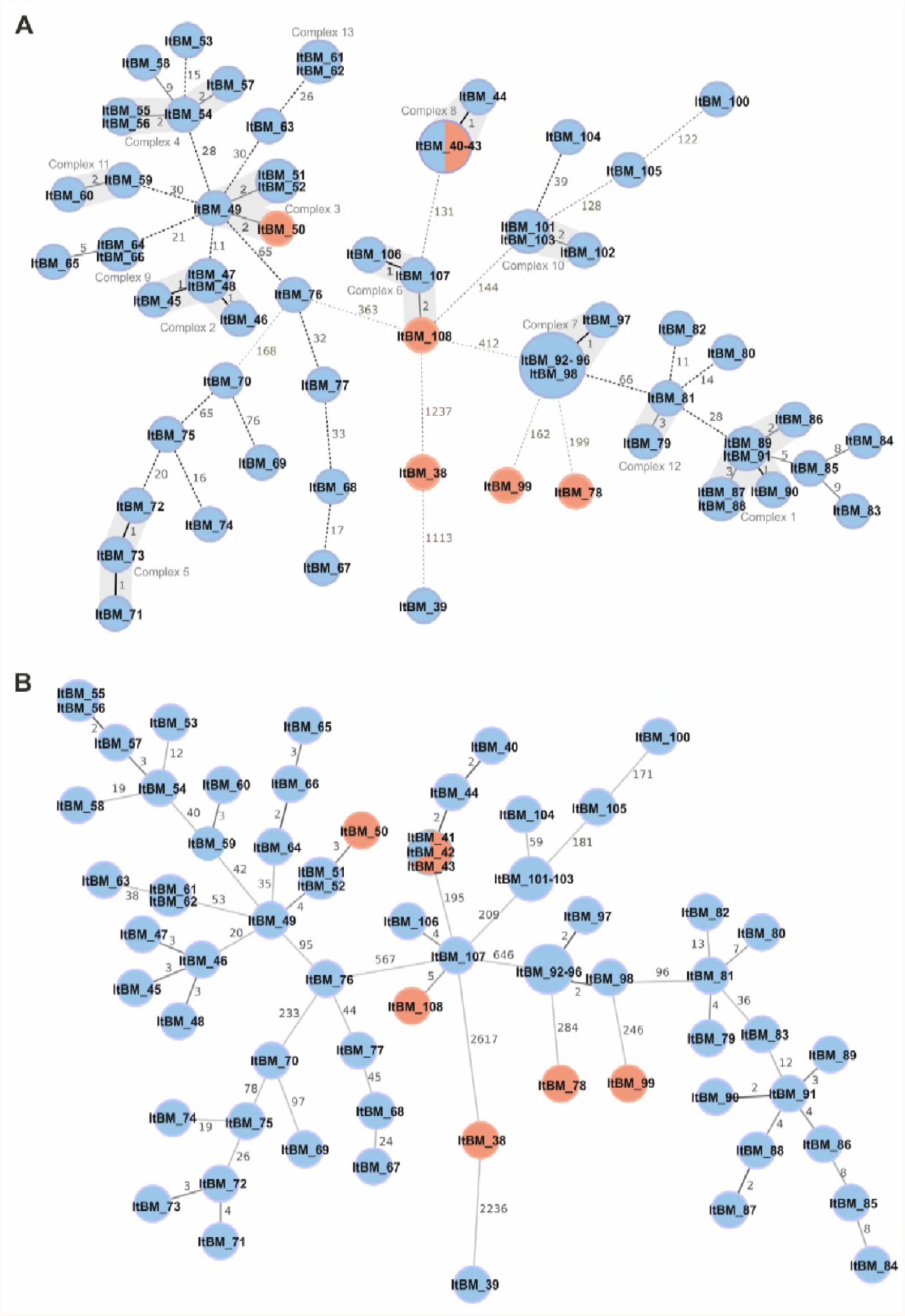
Minimum spanning trees (MST) based on WGS analysis results generated for 71 samples from surveillance epidemiologically unrelated isolates. (A) MST generated using gene-by-gene approach. CgMLST profiles were assigned using *B. melitensis* task template with 2687 target genes. The MST was created by cgMLST target comparison, pairwise ignoring missing values with distance representing the number of diverse alleles. Separate complexes are highlighted. (B) MST based on single nucleotide polymorphism (SNP) analysis using *Brucella melitensis* strain 16M as a reference. The distance labels correspond to the number of discriminating SNPs between neighboring genotypes. The red nodes correspond to human isolates and the blue nodes to animal isolates.

Variation within the West Mediterranean group ranged from 1 to 412 loci. Based on the analysis of outbreak-related isolates we used 4 alleles as a threshold for a potential complex of related cases. Thirteen complexes were therefore assigned in the MST data analysis with Complex 1 containing the highest number of related isolates. Gene by gene analysis confirmed relatedness of genotypes with MLVA profiles 10 and 15, however, according to cgMLST two other isolates were in 0 to 1 gene distance away from the samples of MLVA profile 15, and one other isolate of profile 10. ItBM_55 that was classified as MLVA profile 24 was shown not to be closely linked to other isolates with the same MLVA alleles when examined with a gene by gene approach.

With MLVA typing, only two human isolates were traced back to a probable animal source. Using cgMLST however, four of the human isolates were found in the distance not exceeding 2 alleles to the closest animal strain. ItBM_41 and ItBM_43, originating in Salerno, could be traced to three isolates collected from sheep and goats in the same region (Campania), as seen also in MLVA profiling data. These two samples that were collected from the same patient which was confirmed with the identical cgMLST profile. Another human isolate of *B. melitensis* (ItBM_108) displayed 2-allele variation compared to the closest related animal isolate analyzed in this this study (ItBM_107). Interestingly, the samples were collected two years apart and in different geographical locations suggesting that ItBM_107 could have been closely related (or ancestral) to the source of human infection but not directly involved in the transmission event. A similar scenario was observed for sample ItBM_50 showing a distance of two alleles to an animal isolate. In these cases, observation based on cgMLST typing suggests that strains of *B. melitensis* were circulating in the affected regions of Italy for many years and the surveillance program failed to eradicate them. Two of the human samples originating in Piemonte, although clustered within the West Mediterranean clade (ItBM_99, ItBM_78) were genetically divergent from animal samples, with 162 and 199-allele difference to the closest isolate and could be identified as outliers although distantly related to other Italian genotypes. Epidemiological investigations linked those cases to people who had come from northern African countries where the same West Mediterranean lineage is thought to be prevalent (35). Divergence of these two samples was not however evident in MLVA typing (2-3 alleles of distance to other isolates).

6,533 SNPs were discovered by mapping 71 genomes to the *B. melitensis* 16M reference strain. Out of these, 6,029 were considered high quality discriminatory SNPs and were used to infer the relationship between the strains. Minimum Spanning Tree (MST) analysis using goeBurst function within PHYLOViZ software grouped the isolates based on their SNP profiles (**Figure 4B**). We applied the threshold of 10 SNPs to detect the clusters of closely related cases and, in concordance with cgMLST analysis, we identified thirteen complexes. The highest distance observed between two adjoining isolates were 2,617 and 2,236, belonging to the SNP profiles of ItBM_38 and ItBM_39, which were marked as outliers also by MVLA and cgMLST analyses. Distances of 646 SNPs followed by 567 and 233 SNPs were observed between the West Mediterranean isolates originating in Italy and the tree structure corresponded to the MST based on cgMLST described above.

In agreement with cgMLST, two human cases could not be traced to any of the analyzed animal strains of *B. melitensis* and both differed by more than 200 SNPs from the nearest SNP profile (ItBM_99, ItBM_78), and did not cluster with any of the samples analyzed in this study. Close genetic relationship to one isolate from animal hosts was confirmed for ItBM_41 and ItBM_43. This suggests a close contact of the person with the farm animals (or their products), which resulted in infection with *B.melitensis*. Two human isolates ItBM_108 and ItBM_50, that in previous analyses showed close relationship to strains originating from an animal source, here, were separated only by 5 and 3 SNPs to the neighboring isolates, respectively.

## DISCUSSION

Our study compared the performance of two WGS-based typing methods, SNP analysis and cgMLST with the gold standard MLVA-16 in the analysis of the phylogenetic relationship between isolates of *B. melitensis* detected in the context of the national surveillance program.

We found that all three typing schemes generally performed equally and although SNP analysis had the highest resolving power in terms of differences detected between the isolates, the number of predicted genotypes in the surveillance scenario was comparable for all examined methods (51 MLVA types, 55 cgMLST types, 60 SNP types).

CgMLST accurately predicted the presence of two genomes divergent from the rest of the Italian strains. The distance between each of these and the nearest Italian isolates was shown to be more than 1000 alleles while no more than 412 alleles of difference occurred between any of the local strains. In fact, the majority of analyzed samples belonged to West Mediterranean lineage of *B. melitensis*, while the outliers were members of the East Mediterranean and American lineages (6). The other two methods confirmed this result, with SNP analysis identifying more than 2000 SNPs and 9 MLVA alleles of distance to the closest Italian genotype for both samples. For distantly related genomes from the same lineage, cgMLST as well as SNP analysis provided much higher phylogenetic distance resolution than MLVA, and therefore spotting divergent genotypes, unlikely to be connected to the other circulating strains, was possible with greater confidence. This was particularly apparent in case of two clinical isolates with cgMLST distance of 162 and 199 from the closes animal isolate, and in case of *B. melitensis* collected from an ibex (*Capra ibex* ibex) in National Park Gran Paradiso located in the Graian Alps in Italy (sample ItBM_100). This demonstrated that while all applied schemes could be used to identify very distant genomic outliers within *Brucella* population, WGS-based schemes were superior in identifying unrelated cases belonging to the same lineage. Additionally, within the clusters of similar genotypes, cgMLST performed equally as the SNP analysis, but some discrepancies were observed in MLVA analysis. For instance, seven isolates from Sicily had SNP profiles differing by maximum of two polymorphisms (type 50, 51 and 52) suggesting that they were very closely related. The same result was demonstrated by cgMLST with a difference of only one allele between the genotypes. However, while five of these isolates shared MLVA profile 15, one belonged to type 8 (1-allele distant) and another to type 12 (2-allele distant). The interpretation of WGS results would therefore suggest that these were actually strains from the same cluster, while MLVA typing would not necessary lead to the same conclusion. Similar observation was reported by Dallman *et al*. (2015), who showed that using SNP analysis of *E. coli* O157 isolates identified linked cases with twice the sensitivity of MLVA scheme, while Georgi and colleagues (2017) demonstrated that MLVA had lower discriminatory power than the WGS-based SNP typing by analyzing a set of 63 human *B. melitensis* isolates (36, 37). Interestingly, in our cluster of outbreak-related cases, we identified several genotypes that differed by one or two hypervariable alleles and belonged to the same epidemics. CgMLST analysis of these strains showed that they were very closely related (less than 5 genes of difference). This observation shows that MLVA profiling does not provide enough resolution to discriminate between isolates involved in an ongoing epidemics or strains that have been circulating over the years with no direct link to one another.

SNP analysis has successfully been used to discriminate between *Brucella* species and mapping the geographic distribution and global spread of *B. melitensis* (18, 36, 38). However, to date, there is no official and validated cgMLST or SNP scheme for any of the *Brucella* species. Consequently, the cluster types for specific data, and in particular, for closely related strains, can only be assessed empirically and therefore is subject to variation between the laboratories where the analysis is carried out. In order to reliably interpret the results, cut-off values should be first established based on the analysis of a significant number of closely related strains and unrelated strains sharing common or closely related profiles assigned using gold standard typing methods. In the outbreak-related isolate analysis, a number of variable loci within a cluster did not exceed 4 (cgMLST) or 10 SNPs, suggesting that within 2-year duration of the study, a limited evolution of the strains occurred and the genotypes remained relatively stable. These findings highlight the potential criteria necessary for inclusion of an isolate into a brucellosis outbreak cluster that we would therefore suggest to be ≤4 loci in the cgMLST and ≤10 in WGS SNPs analysis. In surveillance scenario we were able to identify human and animal isolates, collected several years apart, with less than ten alleles of difference between one another in cgMLST scheme. Using our proposed cut-off values, these would not be included in the active outbreak cluster, however the possibility that they shared a common ancestor at some time of their evolution would not necessarily be excluded. In WGS analysis the quality of the reads as well as of the assembly plays a crucial role in achieving reliable cgMLST results. While in our study all samples reached at least 97.9% of good targets when aligned to the task template targets, low quality assemblies are likely to have a reduced number of good targets and therefore lead to generation of inaccurate results in phylogenetic analysis. We would therefore propose that the data with good targets of less than 97% should be taken with caution.

In conclusion, WGS/NGS data can be used effectively to gain a better understanding of epidemiology and dynamics of *Brucella* populations and to gather in depth information which can be used for source tracing in case of outbreaks within animal holdings, zoonotic or food-borne infections and illegal animal movements. Moreover, WGS data facilitates the assessment of the possible extent of an ongoing outbreak and the reliable prediction of the routes of its spread.

In accordance to the *One Health* approach, public health agencies can therefore implement WGS to aid the disease control and eradication plans. In our study, both cgMLST and SNP analysis performed well despite of the restricted level of *B. melitensis* genetic diversity and we demonstrated that the performance of the gene by gene approach was comparable to the SNP analysis. On the basis of these results, we believe that MLVA16 typing of *B. melitensis* in Italy can now be successfully replaced by the more informative WGS analysis.

## ACKNOWLEDGMENTS

This study was funded by the project IZS AM 02/17 RC from the Italian Ministry of Health ricerca corrente 2017 funds, and from the ANHIWA project BruEPIDIA.

## CONFICTS OF INTEREST

D.H. is one of the developers of the Ridom SeqSphere+ software mentioned in the article, which is a development of the company Ridom GmbH (Muenster, Germany) that is partially owned by him. The other authors have declared no conflict of interests.

